# Resourceful mice do not starve: feeding efforts and decision-making process under a restricted unknown food source

**DOI:** 10.1101/546705

**Authors:** M Carmen Hernández, Álvaro Navarro-Castilla, Isabel Barja

## Abstract

Foraging decisions must balance the energy gained, the time investment and the influence of key environmental factors. In our work, we aimed to examine the importance of predation risk cues and experience in the feeding efforts and decision-making process when a novel food resource is presented. To achieve this, free ranging wood mice *Apodemus sylvaticus* were live-trapped in “Monte de Valdelatas” (Madrid) by setting 80 Sherman traps in 4 plots. Traps were subjected to two food access difficulties three-night consecutive treatments: open plastic bottles and closed bottles, both using corn as bait. To generate predation risk, we set fox faeces in half of the traps in each plot. Also, we considered indirect predator cues as the moon phase. We analyse whether mice had bitten the bottles and the area gnawed of each bottle was measured. We discovered that mice feeding decisions and efforts were driven by food access difficulty, experience and predation risk. The ability of mice to properly balance their energy budget was probed since they bit and performed bigger orifices in the closed bottles, hence, individuals can adapt the feeding effort when a new food source is available. Moreover, experience was determinant in the use of this new resource since recaptured mice gnawed the bottles more successfully and the skill was improved each time an individual was recaptured. Additionally, direct predation risk cues prompt mice to bite the bottles whereas the effect of different moon phases varied among the treatments. This is the first study that provides direct evidence of wild mice formidable efficacy to exploit a new nutrient resource while deepening in crucial environmental factors that shape decision-making procedure.

## Introduction

Wild animals must cope with unpredictable environmental demands. In this particular setting, choices made by animals when selecting food and regulating intake aim to satisfy their specific levels of nutrient requirements [1, 2, 3]. The variable time and space food availability challenge animals to select the type of food which best meets their nutrient demands and to evaluate if it counterbalances the energetic effort they have to make to obtain it [4]. These changeable environmental conditions have led to the development of a wide array of adaptations to efficiently satisfy the energetic requirements of all life forms [5, 6, 7], making possible for them to exploit and utilize heterogeneous food sources. The mechanisms which underlay feeding choices are rather diverse, being both endogenous and environmental factors involved in the decision process [8, 9, 10, 11, 12]. It is known that animals possess the ability to learn about the characteristics of the items in their diet and that feeding choices are experience dependent [12, 13, 14, 15, 16]. In this manner, experience and learning can provide animals the key to quickly adapt to this ever-changing environment by displaying novel feeding strategies when new food sources are present.

On the other hand, there is persuasive evidence of predation risk influence on prey’s behaviour [16, 17, 18, 19, 20, 21], complicating the decision-making process even more when it comes to feeding opportunities. Prey animals possess the ability to estimate predation risk and adjust their behaviour to reduce the probability of being preyed [22, 23, 24], which is critical in habitats where the magnitude of threats is spatially and temporally mutable [25, 26, 27]. Chemosensory cues are of vital importance for predation risk assessment in mice [16, 28]. These chemical signals are crucial for prey species since it can alert them of the presence of any potential predators and procure information about their activity and diet [29], modulating daily activity patterns [21, 30, 31] and feeding habits of preys [32]. Moreover, perceived predation risk can vary depending on environmental factors such as habitat complexity and moonlight [21, 33, 34]. The influence of moonlight on mammal’s behaviour and its relationship with predator-prey dynamics is well documented [35, 36, 37, 38, 39]. For rodents, bright nights increase detectability by predators and hence, predation risk. As a consequence, rodent species tend to decrease their activity near to full moon nights [20, 40, 41, 42, 43]. Hence, for prey species, feeding strategies should be a trade-off between predation risk avoidance and the benefits of obtaining energy [19, 20, 44, 45, 46]. However, behaviours that maximize food intake often increases exposure to predation risk, so preys must gather all the environmental information, decide how to allocate resources and pursue the option which maximizes their fitness [47]. Therefore, properly balancing the energy budget should be an important selective force for the evolution of life-history traits.

The aim of this study was to analyse feeding efforts under restricted food access conditions in the wood mouse (*Apodemus sylvaticus*). Concretely, we focused on studying mice feeding behaviour when facing a new food resource with limited access and unravelling the importance of experience testing the ability to learn and develop new effective strategies in a brief period of time to maximize food obtaining. Furthermore, we also evaluated if feeding efforts performed under different food access restriction were conditioned by predation risk cues (predator faeces and moonlight). On one hand, we predicted that mice feeding efforts would be certainly influenced by the difficulty of the food access. We expected that individuals only would spend energy trying to gain access to food if it is necessary. Thus, mice facing an easier food access restriction should spend less energy trying to reach the bait than those ones facing a more complicated food access treatment. On the other hand, it was expected that recaptured individuals would have developed a more efficient feeding technique, allowing them accessing food in an easier way than those ones which do not have previous experience with this kind of food resources. Finally, we also expected diminished food efforts in those traps treated with fox faeces and during brighter nights, due to a higher perceived predation risk causes a decrease in the activity of mice [20].

## Materials and methods

### Study area

The research was conducted in the “Monte de Valdelatas” (Madrid, Spain), a Mediterranean forest located at an altitude of 650 m a.s.l. The characteristic vegetation is forests of holm oak (*Quercus ilex ballota*) and scrubland (gum rock roses *Cistus ladanifer*, thyme *Thymus zygis* and umbel-flowered sun roses *Halimium umbellatum*). Wild predators are frequent in this habitat, being of importance the red fox (*Vulpes Vulpes*) and the common genet (*Genetta genetta*) [19, 48].

### Live-trapping and data collection

Fieldwork was performed in March 2017-2018 in four plots with similar vegetation and composition. The distance between plots was 35 m to ensure that they were independent and that they corresponded to different mice populations [16, 28]. In each plot, 20 Sherman^®^ live traps were set in in a 4 x 5 grid with 7 m of distance among them [16, 28]. Total trapping effort was 960 traps-night (20 traps x 4 plots x 3 nights x 2 food treatments x 2 trapping sessions). All traps were hidden under vegetation cover to protect animals from adverse weather conditions and bait was provided inside traps (see details below). Traps were opened at sunset and data collection was daily started after the sunrise.

All captured animals were identified to species by external morphology and each captured mouse was weighed with a scale (PESNET, 100 g, PESNET 60g). Sex and breeding condition were checked according to Gurnell and Flowerdew [46]. Sex was determined using the anal-genital distance, which is longer in males than in females. In breeding adult males, the testicles were bigger, whereas breeding adult females showed conspicuous nipples in the abdomen and thorax and the vaginal membrane appeared perforated. Harmless waterproof paints (Marking stick DFV, EXT-link: www.divasa-farmavic.com) were used to mark captured individuals in non-conspicuous areas (e.g. ears, toes and tail) for discriminating recaptures [49]. Finally, all captured animals were immediately released after handling in the same place of capture.

### Predation risk simulation

To simulate predation risk, we used red fox faeces since this species is known to be present in the study area [19, 48] being one of the most common small mammal predators [50, 51]. Furthermore, red fox faeces have been previously demonstrated to elicit antipredatory responses effectively [19, 20, 28, 52]. Fresh faeces used for the treatment were obtained from captive red foxes (one male and one female) on a carnivorous diet from the Centro de Naturaleza Opennature Cañada Real (Peralejo, Madrid). We considered as fresh faeces only those ones with a layer of mucus, an elevated level of hydration and strong odour [53, 54], and all faecal samples were frozen at −20 ºC until treatment preparation. Seasonal and individual factors are known to influence volatile compounds variation among individuals [55, 56, 57, 58] so, to guarantee homogenization (providing a similar degree of predation risk in all the treated traps, and therefore) and avoiding possible result bias, all collected red fox faeces were properly mixed.

In each plot, half of the traps were subjected to a predator odour treatment consisting in 2 g of fresh fox faeces. Within the 4×5 grids set in each plot, predator treatment was set on two non-consecutive rows (10 traps) while the other two rows (10 traps) acted as controls (i.e. without predator faecal cues). In order to avoid the influence of border effects due to treatment distribution, control and predator treatment rows were alternated in each plot. The faecal material was placed on one side of the trap entrance to avoid blocking the entry for rodents but close enough to act as a potential predation risk cue (i.e. 3 cm approximately). Predator treatment was replaced every day at sunset to guarantee odour effectiveness when mice are more active, i.e. two or four hours after the dusk [59].

Regarding indirect predation risk cues, since mice are known to be more active when moonlight is dim due to a reduced predation risk perception [20, 40, 41, 60], we avoided trapping during high illuminated conditions (i.e. full moon phase and closer nights). Thus, live-trapping sessions were carried out under low (< 25%, new moon) and medium (25-54%, waxing crescent phase to the beginning of the first quarter) moonlight conditions. Moon percent illumination corresponding to each sampling night was downloaded from the AEMet website (National Meteorological Service, www.opendata.aemet.es).

### Food access experiments

All traps were subjected to two different consecutive food access treatments in which food access difficulty was experimentally manipulated using polyurethane plastic bottles of 6 cm length, 2,7 cm of total diameter and 2 cm of aperture diameter, baited with 5 g of toasted corn inside. First treatment (first three nights) consisted in opened plastic bottles inside all traps while for the second treatment (next three consecutive nights) all traps were provided with baited closed bottles (we performed ten 1 mm holes with a needle in order to allow mice to smell the bait).

After trapping sessions, plastic bottles from the experiments were analysed in the laboratory to determine mice feeding efforts. For each bottle, we firstly confirmed mice handling through the presence or the absence of bite marks made by individuals. To quantify feeding efforts, we measured the total area gnawed by each mouse (i.e. size of the orifice performed in the bottle). For this, gnawed areas were exactly transferred to translucent paper sheets and they were scanned. Later, to measure the gnawed area, we analysed the scanned sheets through the Adobe Photoshop CC® software in a similar way to [61], selecting the target gnawed area with the *magic wand* tool and using the image analysis tool to know the gnawed area size in pixels.

Finally, to determine the amount of food eaten by each individual, we collected the unconsumed bait from each trap. The remnant bait was dried at 80 ºC in a heater for 1 h to eliminate moisture and weighed with an electronic balance (C-3000/0.01 g CS, COBOS; precision 0.01 g). Thus, food intake by each individual was obtained by deducting the remnant bait weight to the initial 5 g of corn supplied inside each bottle.

### Statistical analysis

Since model residuals were not normally distributed, behavioural responses were analysed using Generalized Linear Models (GLMs). Robust estimator (Huber/White/ sandwich estimator) was used to correct homogeneous variances criteria deviations. To analyse factors triggering mice handling of plastic bottles we performed a binomial distribution logit link GLM being the response variable the presence or absence of bite marks in the plastic bottles. Furthermore, to assess feeding effort, we use a GLM with normal distribution and identity link, being the response variable the missing area gnawed by mice in each bottle measured in pixels. For both models, the explanatory variables considered were the same: food access (opened bottle/closed bottle), recapture (first captured/recaptured), moonlight (new moon/waxing crescent), predation risk (control/predator), reproductive status (breeding/non-breeding) and sex (female/male), including weight as a covariate. We also tested the interactions food access*recapture and food access*moonlight. Furthermore, we also conducted separate ANOVA tests to analyse whether the gnawed area varied through repeated consecutive recaptures. Finally, a nonparametric Spearman's correlation analysis was performed to check the relationship between the effort made by mice to obtain the bait (gnawed area) and food intake. Because mice did not need to gnaw open bottles to obtain the bait provided and due to the statistically significant relationship between food access with the extension of the gnawed area by mice, we only considered data from closed bottles for this correlation analysis.

Results were considered significant at α < 0.05. Data are represented as mean ± standard error (SE). The software used to perform the statistical analysis was SPSS 23.0 for Windows (SPSS Inc, Chicago, IL, USA).

## Results

The total number of captures was 142, corresponding to 84 different individuals. Results of the binomial model showed that food access, recapture, predation risk and the interaction between food access and moonlight were the factors which explained the presence of bite marks in bottles (Table 1).

**Table 1.** Results of the binomial logit GLM analysing the effect of individual, environmental and experimental factors on the absence or presence of bite marks performed by mice in the plastic bottles.

| Factor | *F* | df | *p* |
| --- | --- | --- | --- |
| Food access | 14.113 | 1 | 0.000 |
| Recapture | 7.618 | 1 | 0.006 |
| Moonlight | 1.772 | 1 | 0.183 |
| Predation risk | 5.945 | 1 | 0.015 |
| Reproductive status | 0.022 | 1 | 0.883 |
| Sex | 2.627 | 1 | 0.105 |
| Weight | 0.242 | 1 | 0.623 |
| Food access*Recapture | 0.049 | 1 | 0.826 |
| Food access*Moonlight | 4.017 | 1 | 0.045 |

In open bottles (N= 89), only the 33.7% showed bite marks whereas in the closed bottles treatment (N= 53) the 90.6% of them were bitten by mice. The 75.9% of the recaptured mice bitted bottles (N= 58), while this percentage decreases to 40.5% for first-captured ones (N= 84). As for the predation risk influence, we found bite marks in 67.5% (N= 51) of the bottles treated with fox faeces, being this percentage lower in the absence of predator cues (50.0%, N= 27). Regarding the interaction between food access and moonlight, we found that mice bite marks were particularly less frequently found in open bottles during new moon nights (27.8%, N=20), while this percentage was higher during waxing crescent (58.8%, N=10). By contrast, bite marks appeared in the majority of the closed bottles independently of the moon phase: new moon nights 95.7% (N= 22) and 86.7% (N= 26) during waxing crescent nights.

Results of the GLM analysing mice feeding efforts (i.e. gnawed area) are showed in Table 2; main influencing factors were food access, recapture and moonlight. The average area gnawed by mice in open bottles was lower (6690.0 pixels ± 2141.0 SE) than in closed ones (26277.4 ± 4361.0). Overall, recaptured individuals gnawed an average area of 24864.3 ± 4090.5 pixels, while a reduced area of 6499.8 ± 2213.9 was performed by first-captured mice. Interestingly, separate analyses showed that the area gnawed by mice exponentially increased during consecutive recaptures (*F*_4,48_= 7.641, p< 0.001), but this significant effect was driven by individuals facing closed bottles (*F*_4,48_= 3.226, p< 0.05) (Fig. 1).

**Table 2.**
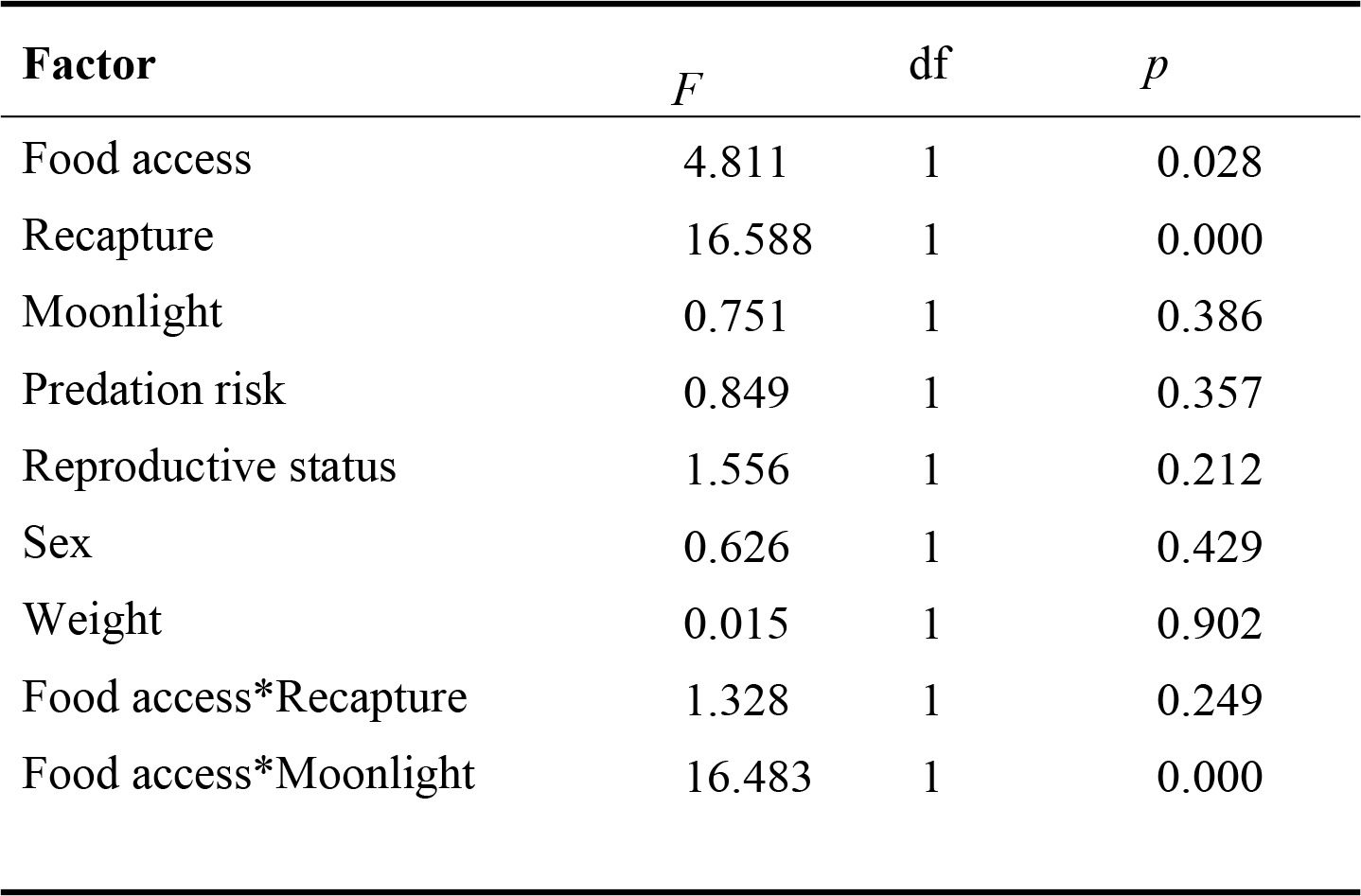
Results of the GLM testing the effect of individual, environmental and experimental factors on feeding effort (area gnawed by mice).

| **Factor** | *F* | df | *p* |
| --- | --- | --- | --- |
| Food access | 4.811 | 1 | 0.028 |
| Recapture | 16.588 | 1 | 0.000 |
| Moonlight | 0.751 | 1 | 0.386 |
| Predation risk | 0.849 | 1 | 0.357 |
| Reproductive status | 1.556 | 1 | 0.212 |
| Sex | 0.626 | 1 | 0.429 |
| Weight | 0.015 | 1 | 0.902 |
| Food access*Recapture | 1.328 | 1 | 0.249 |
| Food access*Moonlight | 16.483 | 1 | 0.000 |

**Figure 1.**
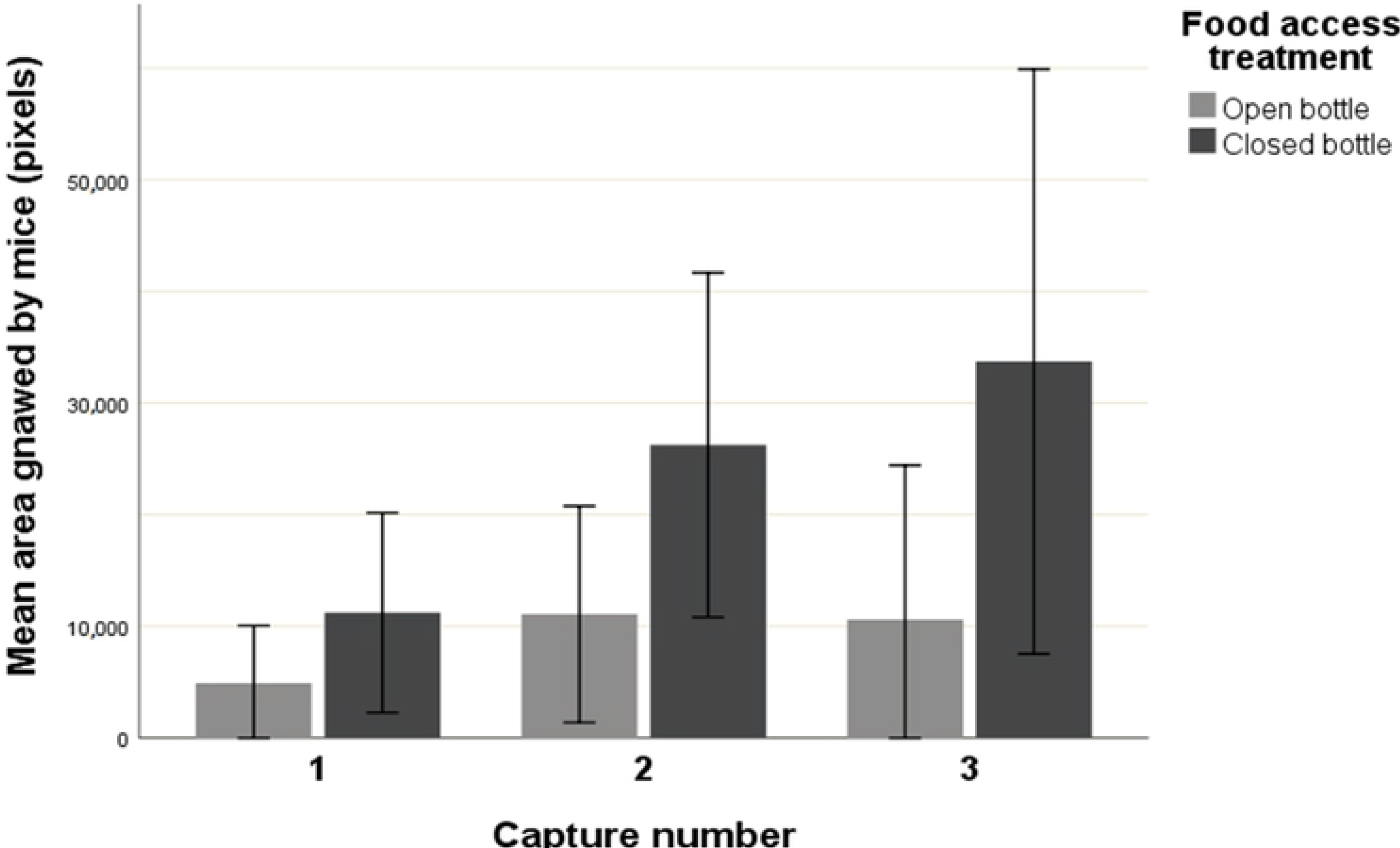
Mice feeding effort (mean area gnawed ± SE) through consecutive captures of each individual depending on the food access treatment (open bottle / closed bottle).

Furthermore, the interaction between food access and moonlight showed that mice gnawed particularly broad areas in the closed bottles during new moon nights (45373.4 ± 7735.7) (Fig. 2). Finally, a correlation analysis showed that there was a positive correlation between the effort made (i.e. area gnawed) to obtain the bait and mice food intake (Spearman correlation, r= 0.805, N= 142, p < 0.0001).

**Figure 2.**
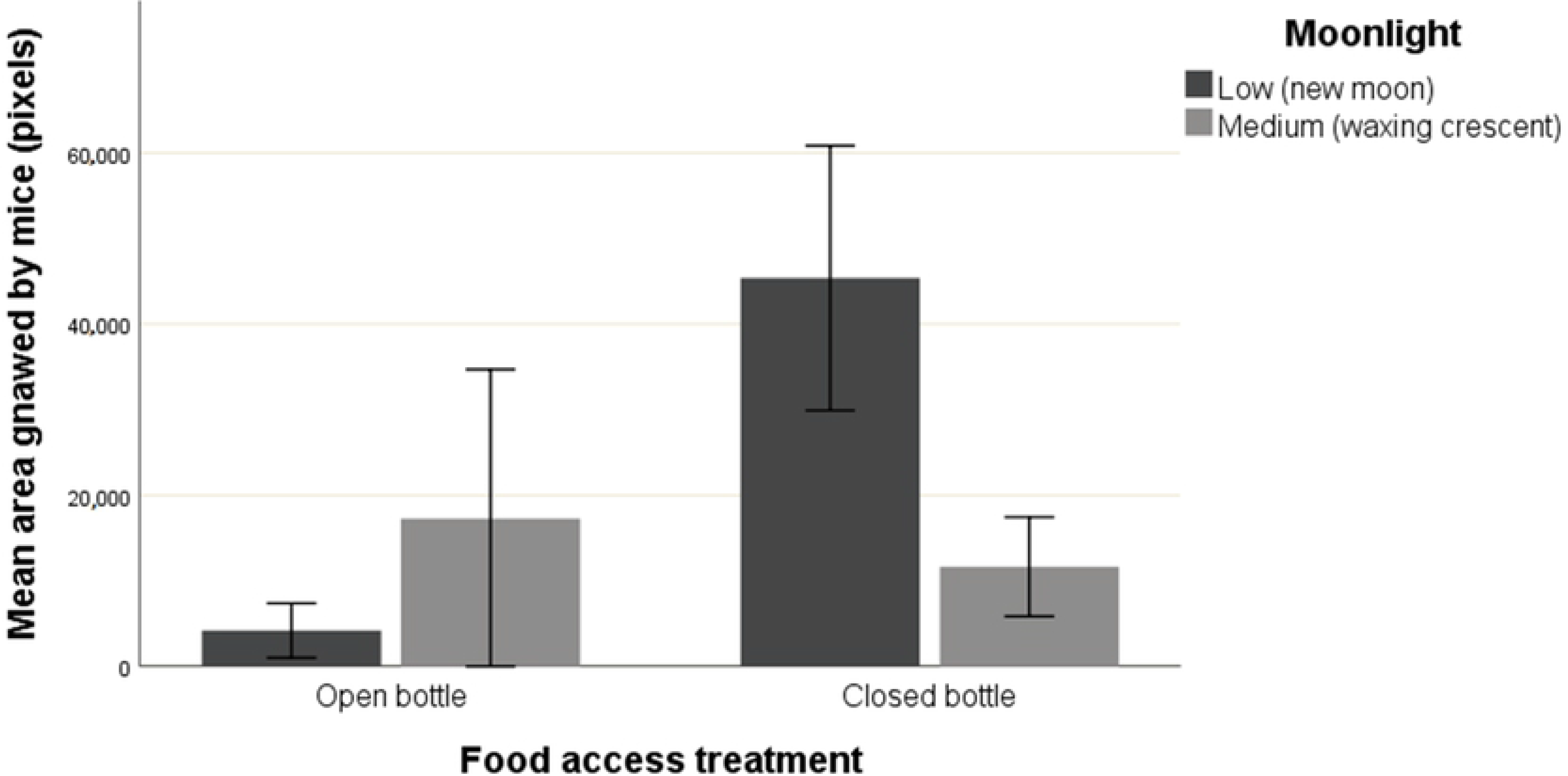
Mice feeding efforts (mean area gnawed ± SE) in relation to food access (opened bottle or closed bottle) and moonlight (low, new moon / medium, waxing crescent).

## Discussion

To our knowledge, this is the first study which provides evidence of the importance of experience and perceived predation risk in wood mice feeding efforts and decision-making process. As expected, food access difficulty determined the presence of bite marks in the bottles, probing that mice understood the implications of the feeding devices since they tended to spend extra energy on food handling only if it was mandatory (i.e. closed bottles). Moreover, experience also determined mice choices in relation to bite or not to bite the food container. Naïve individuals were less inclined to gnaw the plastic bottles, demonstrating that experience is a decisive factor regulating wood mice feeding choices when a new source of food is available [12]. Predator cues also affected mice decision-making process, in this case, fox chemical signals seem to have a stimulating effect which prompted individuals to interact with the food containers. Predator scents have been previously demonstrated to modify food intake [16, 17, 19, 62], however, the direction of this association is not clear since there is evidence of both a rise and a decrease in the food intake. In our study, we hypothesise that traps could have provided mice a safe space to handle the food resources [16, 63], as a consequence, mice might have chosen to feed because they were sheltered against predator attacks. Alternatively, predation risk could have trigger physiological stress response in mice [28] and the immediate mobilization of energy could have stimulated mice to bite the food containers.

Regarding the food access and moonlight interaction effect, while mice facing open bottles were more reluctant to try to get access to food during new moon nights, the moonlight did not influence mice behaviour when bottles were closed. When experience closed bottles, mice are compelled to bite the containers to obtain the food in spite of predation risk cues. In this particular setting, the prospect of obtaining a potentially highly nutritious food could counterbalance the risk of being detected [64, 65]. On the other hand, when biting the food containers is not required to accomplish feeding, individuals behave different depending on indirect predator cues. During new moon nights, prey success to detect predators and competitors could be affected [66, 67], thus, to be prepared to display fight or flight responses and to avoid unwanted interspecific interactions, mice could have decided to be more cautious and to save energy to cope with unpredictable events [32].

As for the feeding effort, in accordance with the previous result, food access difficulty determined the extent of mice feeding endeavour, demonstrating that individuals adaptively adjust their energy expenditure depending on food accessibility and avoid to waste energy. Experience and learning have proved to be excellent adaptive features when it comes to feeding [68, 69, 70, 71, 72], making individuals extremely resourceful and giving them the essential responses to survive in highly variable environments. Our study showed that experience prompted individuals to invest energy trying to gain food access and the skill of the procedure was more efficient, since they managed to perforate a wider area of the bottles. In addition, the positive correlation found between the gnawed area and food intake, confirm that the endeavour they performed was justified, spending more energy only if they can counterbalance the feeding costs associated [73, 74, 75]. Our results indicate that mice are fast learners, improving their skill twofold with only a single previous encounter with the food containers. However, this endeavour was only significantly improved in mice facing closed bottles, demonstrating again the ability of individuals to make efficient energy budget decisions. The relevance of experience and learning upon mice feeding efforts is clear, providing mice the opportunity to exploit new food resources in a relatively short amount of time. Despite learning feeding techniques can have expensive associated costs in terms of energy and time [69], the highly variable natural living conditions could have induced the development of this remarkable evolutionary strategy by enhancing mice individual fitness [11, 76].

As for the influence of the interaction between food access and moonlight on feeding effort, new moon nights were associated with increased feeding efforts when individuals were dealing with the more arduous treatment (i.e. closed bottles). This result gives us direct insight of mice decision-process and the behavioural response elicited when a trade-off between predation risk and feeding is presented (see predation risk allocation hypothesis [77]). According to this theory, individuals would increment feeding effort during new moon phase when perceived predation risk is low, since moonlight can increase prey detectability and hence, hunting success for predators [78, 79]. Thus, darker nights caused mice to feel safer, allowing individuals to spend energy in the device handling costs. On the contrary, a rise in perceived predation risk caused by the increase in the moonlight probably caused mice to keep a low profile and to choose survival over increasing their exposure handling the food resource, even though the energetic reward was high. Further, this result would be in accordance with previous studies that show how mice activity and food intake diminish with the increase in night luminosity [20, 42, 43). On the other hand, for opened bottle treatment, the feeding effort remained low during both new moon and waxing crescent because it was not necessary to perforate the bottle to obtain the food, thus, it would be expected that animals did not spend energy when it was not required.

Contrary to our predictions, predator faecal cues did not affect mice feeding efforts. Nevertheless, this result would be in accordance with other studies that discovered no effect of predator cues on feeding behaviour [20, 21, 80]. As we suggested before, traps could have been perceived as a refuge against predators, allowing them to feed in a secure environment [16, 63]. Another plausible explanation would be that due to individuals remained several hours under the influence of this predation cues, they have to resume their feeding activity in order to not compromise their survival [77, 81].

Additionally, we found that individual variables, such us breeding condition, sex or weight, had no effect on feeding behaviour. It could be possible that the higher energetic demands of certain individuals were only reflected upon the food intake rather than having an influence on mice feeding efforts. Although this was not expected, the results clearly show that these factors were not determinant, and that experience and moonlight were the phenomena which modulated wood mice feeding choices and efforts when a new source of food is available. The wood mouse plays a key role in the ecosystems, being a pivotal part of the diet of many often endangered predators [82, 83, 84, 85]. These results provide certain hope about the resilience and plasticity of mice populations, frequently subjected to human-induced changes that can modify food resources and its availability.

